# Sex Differences in Affective States and Association with Voluntary Ethanol Intake in Sprague Dawley Rats

**DOI:** 10.1101/2021.12.02.470921

**Authors:** SG Quadir, GM Arleth, JV Jahad, M Echeveste Sanchez, MA Herman

## Abstract

Alcohol use disorders (AUDs) are a major problem across the United States. While AUD remains a complex human condition, it is difficult to isolate the directionality of anxiety and ethanol (EtOH) drinking from outside influences. The present study sought to investigate the relationship between affective states and EtOH intake using male and female Sprague Dawley rats. Using complementary tests of anxiety- and depressive-like behavior, we found sex- and test-specific differences in basal affective behavior such that females displayed enhanced anxiety-like behavior in the Splash Test and males displayed enhanced anxiety-like behavior in the Novelty Suppressed Feeding Test. Although there were no sex differences in EtOH intake and no correlation between anxiety-like behavior and subsequent EtOH intake, we did find that depressive-like behavior predicted future EtOH intake in females rats only. In addition, we observed an increase in depressive-like behavior is male rats in both the water and EtOH drinking groups. Furthermore, anxiety-like behavior, but not depressive-like behavior predicted subsequent EtOH intake in female rats. Lastly, we found a history of EtOH intake decreased pain thresholds in male and female rats. Together, these experiments provide important information on the complex interaction between negative affect and alcohol intake and how these two contexts reciprocally do, or do not, influence each other in a sex-specific manner.

## INTRODUCTION

Alcohol use disorders (AUDs) are a major problem across the United States. In 2019, studies found over 14 million adults in the United States met the criteria for an AUD (2021). A person’s risk for AUD depends on a number of factors, including age of first drink, familial history of AUD, and mental health disorders (Yuen et al., 2020, Kendler et al., 2020, Kushner et al., 2005). Specifically, AUDs are often comorbid with affective disorders such as anxiety and depression (Kushner et al., 2005). Indeed, one study found that 46% of individuals with AUD also met the criteria for generalized anxiety disorder (Smith and Book, 2010). Similarly, 33% of individuals with AUD also met the criteria for major depressive disorder (McHugh and Weiss, 2019). Not only are these comorbidities extremely common, they can often result in significantly worse outcomes for the individual. Indeed, individuals with comorbid affective disorders and AUD are less likely to seek treatment and even in treatment centers, report increased rates of relapse (Driessen et al., 2001, Schellekens et al., 2015, Schneier et al., 2010, Haver and Gjestad, 2005).

There are currently three theories proposed to explain the prevalence of AUD/affective disorder comorbidities: 1) the anxiety pre-dates the alcohol use, and individuals with anxiety are using alcohol to self-medicate, 2) anxiety manifests as a result of heavy alcohol use, and 3) there is a third factor involved, such as age and developmental experience, (Smith and Randall, 2012). There is a wealth of evidence to support the idea that in some cases anxiety pre-dates alcohol use, where individuals with higher anxiety levels drink more alcohol than individuals with less severe anxiety (Menary et al., 2011). Indeed, 50% of individuals receiving treatment for AUD reported having a baseline anxiety disorder (Kushner et al., 2005). Additionally, individuals with baseline anxiety report significantly more alcohol withdrawal symptoms (Kushner et al., 2005) and are more likely to relapse compared to those without basal anxiety (Kushner et al., 2005). Furthermore, individuals who report using alcohol to self-medicate for their anxiety/depression display increased rates of developing and maintaining an AUD (Crum et al., 2013, Menary et al., 2011, Driessen et al., 2001). However, if anxiety was a primary motivating factor in alcohol use, then treating anxiety symptoms should decrease alcohol intake, which this has not been the case. One study investigated the effects of paroxetine (a treatment for social anxiety) on alcohol use and found no effect of paroxetine treatment on quantity and frequency of drinking days (Thomas et al., 2008). The effect of alcohol use on anxiety has also been extensively studied. Commonly referred to as hyperkatifeia, or the dark side of addiction, this theory states that as the euphoric effects of alcohol wear off, dysphoria ensues as a result of increased stress and decreased reward system signaling (Koob and Volkow, 2016). Indeed, many individuals report increased emotional distress and anxiety following excessive alcohol intake (McKinney and Coyle, 2006, Eriksson et al., 2020) and the severity of emotional distress is directly correlated with amount of alcohol consumed (Van de Loo et al., 2017), suggesting that alcohol dose-dependently induces a negative emotional state during withdrawal. However, the potential that alcohol-anxiety comorbidities are due to a third factor, such as age, are also yet to be resolved. For example, individuals who begin drinking before the age of 15 are 5x more likely to develop an AUD (2021). However, individuals with baseline anxiety disorders also begin drinking earlier (and transition to AUD earlier), thus the role of age and its interactions with anxiety and AUD remain unclear (Kushner et al., 2011).

Another major factor often overlooked in both affective disorders and AUD is sex. The lifetime prevalence of anxiety disorders is 60% higher in women as compared to men (Donner and Lowry, 2013). Similarly, women are 1.71x more likely to develop depression compared to men (2021, Gater et al., 1998, Weissman et al., 1996). While AUD is more prevalent in men (Grant et al., 2015), the gap has been closing: from 2000-2013, the prevalence of AUD in women increased 84% while only increasing 35% in men (Grant et al., 2017). Women also present with more severe AUD and are less likely to enter treatment compared to men (Holzhauer et al., 2020). In addition, despite AUD rates being higher in men, women are 1.75x more likely to develop comorbid AUD and anxiety (Kessler et al., 2005). While sex differences in AUD and affective disorders have been observed for more than 20 years, a majority of the research conducted in both humans and rodents has only focused on males (Guizzetti et al., 2016).

While AUD remains a complex human condition, it is difficult to isolate the directionality of anxiety and alcohol drinking from other influences such as age of alcohol exposure, prior trauma, and/or other environmental factors. However, using preclinical rodent models, we are able to isolate 1-2 variables at a time – in our case, sex and alcohol exposure, and determine directional associations between alcohol and negative affect. The present study sought to investigate the relationship between affective states and alcohol intake using male and female Sprague Dawley rats. Rats were first tested for basal anxiety- and depressive-like behavior and then subjected to 4-5 weeks of voluntary drinking. During this time, the rats were reassessed for anxiety- and depressive-like behavior, as well as for thermal sensitivity. Correlational analyses were performed to determine the influence of basal affective state on future alcohol intake, as well as the potential dose-dependent relationships between alcohol consumption and subsequent affective behavior.

## METHODS

### Subjects

64 Sprague-Dawley rats (32 male, 32 female; 200-250g upon arrival) were ordered from Envigo (Indianapolis, IN). Rats were single-housed in a temperature- and humidity-controlled vivarium under a 12h light/dark cycle (lights on: 07:00 AM; lights off: 07:00 PM) and maintained on Irradiated PicoLab Select Rodent Diet 50 IF/6F Diet (LabDiet, St Louis, MO). Unless otherwise noted, rats were provided food and water *ad libitum*. All procedures adhered to the University of North Carolina Chapel Hill’s Institutional Animal Care and Use Committee. All behavioral testing occurred during the light cycle, and the order of rats alternated between male and female to control for time of day.

### Splash Test

Rats were habituated to the testing room for at least 1h before testing. At the start of each test session, the rat was sprayed with 10% w/v sucrose (spray distance ≈ 5cm) on the dorsal coat before being placed into the apparatus and video-recorded for 10 min, as previously described (Calpe-Lopez et al., 2019, Machado et al., 2012, Butelman et al., 2019, Okhuarobo et al., 2020, Sampedro-Piquero et al., 2020).

The apparatus consisted of a clear plexiglass box measuring 50cm (*l*) x 50cm(*w*) x 38 cm (*h*). The cabinet walls were covered in black and white horizontally striped cardboard and only red light was provided in the apparatus.

Videos were scored using the Behavioral Observation Research Interactive Software (BORIS) program (Friard and Gamba, 2016). Latency to groom, total time spent grooming and time spent rearing were recorded by an experimenter blind to the experimental status of each animal. 1 animal was excluded from splash test analysis because of lighting malfunction during testing.

### Novelty Suppressed Feeding (NSF)

48h prior to the novelty suppressed feeding test, each rat was given 1 Froot Loop (Kellogg’s) in the homecage to reduce neophobia during testing. 24h later, all animals underwent 23h food deprivation. The rats were habituated to the testing room for at least 1h before testing. This procedure was carried out according to previously-established protocols, consistent with previous studies (Garcia-Garcia et al., 2016, Sidhu et al., 2018, Jury et al., 2017, Torruella-Suarez et al., 2020). Briefly, the rat was placed in the brightly lit (150 lux) testing apparatus (50cm (*l*) x 76cm (*w*) x 40cm (*h*)) with a single Froot Loop sitting on a piece of filter paper (diameter: 7cm) in the center of the apparatus and latency to begin eating the Froot Loop was recorded. The test was terminated when the rat initiated feeding of the Froot Loop, or after 10 minutes had transpired since being placed in the test apparatus. Upon termination of the test, the rat was placed back in its home cage with a pre-weighed amount of Froot Loops (≈ 8 Froot Loops). The rat was then left undisturbed for 5min, upon which the experimenter removed the Froot Loops and regular chow was returned. The total amount of Froot Loops consumed during this 5 min post-test was normalized to body weight.

### Intermittent Access to 2 Bottle Choice (IA2BC)

After all baseline testing was completed, rats received 2 water bottles on the homecage to accustom them to drinking out of sipper tubes for 5 days before start of IA2BC. Rats were randomly split into ethanol (EtOH) and water drinking groups. Rats in the ethanol group received 2 bottles: one containing 20% v/v EtOH, and one containing filtered water. Consistent with other studies, the bottles were left on for 24h on alternating days (Monday, Wednesday, Friday: bottles on at 10am; Tuesday, Thursday, Saturday: bottles off at 10am; Sunday: undisturbed) (Quadir et al., 2020b, Quadir et al., 2020a, Hwa et al., 2011, Wise, 1973, Darcq et al., 2016, Barak et al., 2011a, Barak et al., 2011b, Carnicella et al., 2009, Carnicella et al., 2008, Simms et al., 2008, Priddy et al., 2017). After 24h, EtOH and water bottles were removed, rinsed, and refilled with filtered water for the days off. Placement of the EtOH bottle (left vs right) was alternated on drinking days to account for any side bias. Animals in the water group had both of their bottles removed and replaced every other day to ensure each cage was handled the same way. Beginning at the fifth week of EtOH drinking, bottles were briefly removed (~30s) at the 1h timepoint for measurements. This timepoint investigates the volume that rats drink immediately when EtOH is presented as compared to drinking that is spread out throughout the 24h access period. Body weights for all animals were taken once weekly on Monday (before bottles went on) and used for that week’s measurements. 2 cages without animals were used to account for bottle spillage, and the average spillage per day was subtracted from each animal’s intake before normalizing intake to body weight.

### Thermal Sensitivity

Thermal sensitivity was assessed using the heated Plantar Analgesia Meter (IITC, Woodland Hills, CA), similar to other studies (Saika et al., 2015, Cheah et al., 2016, Tsiklauri et al., 2017). The rat was habituated to the heated glass surface for 50 min before testing began. After habituation, the experimenter blind to the animal’s condition shined an infrared light (artificial intensity: 40) onto each hind-paw and recorded the latency to withdraw. This was repeated twice for each paw, unless the latencies differed by >1s, in which case a third measurement was taken. Measurements for each paw were averaged and then the latencies for each paw were averaged together in order to obtain a final latency time. A cut-off time of 20s was used to prevent tissue damage. All testing occurred during the light cycle.

### Statistics

All statistical analysis was performed using Graphpad Prism 8.0 (San Diego, CA). In all studies, the threshold for significance was set to *p*<0.05. For baseline Splash Test and NSF, an unpaired Student’s *t* test was used. For drinking studies, a two-way mixed ANOVA was used with time as within-subjects and sex as a between-subjects factor. For correlational analyses, Pearson’s Correlation test was used. For the post-drinking splash test, a two-way mixed ANOVA was used (Pre-test vs Post-Test: Within, EtOH drinking: Between). To assess thermal sensitivity, a two-way between subjects ANOVA was used (factors: sex, EtOH drinking). When applicable, Bonferroni’s test was used for post hoc analysis.

## RESULTS

### Baseline Affective State

To investigate potential sex differences in anxiety- and depressive-like behavior, male and female rats underwent behavioral tests before and after chronic drinking as illustrated in the experimental timeline in **Fig 1A**. The rats first underwent the splash test (**Fig 1B**) in which the latency to groom, total time spent grooming, and time spent rearing were used as measures of anxiety-like, depressive-like, and exploratory behavior, respectively. In the splash test, females exhibited an increased latency to groom compared to males (unpaired Student’s t test: t=2.814, df=62, p<0.001, **Fig 1C**) but there were no sex differences in the total time spent grooming (unpaired Student’s t test: t=0.161, df=62, **Fig 1E**). In addition, females spent more time rearing as compared to males (unpaired student’s t test: t=2.868, df=62, p<0.01, **Fig 1E**). Anxiety-like behavior was further examined through the Novelty Suppressed Feeding test (NSF, **Fig 1F**). In this test, latency to eat is used as a measure of anxiety-like behavior, and post-test consumption is used as a measure of overall motivation to eat. In contrast to what was observed in the splash test, females exhibited decreased latency to eat as compared to males (unpaired Student’s t test: t=2.454, df=62, p<0.05, **Fig 1G**) but there were no sex differences in post-test consumption (unpaired t test: t=0.559, df=62, **Fig 1H**), indicating that the differences in latency were not confounded by differences in motivation to eat. As latency to groom (splash test) and latency to eat (NSF) are commonly-used tests of anxiety-like behavior, we examined the correlation of these two behaviors. Interestingly, both sexes demonstrate a positive but not significant correlation (Pearson’s r_males_(30)=0.134, Pearson’s r_females_(30)=0.319, **Fig 1I**). Together, these data suggest that there are baseline sex differences in anxiety- and depressive-like behavior, but that these differences are test and/or modality specific.

**Figure 1.**
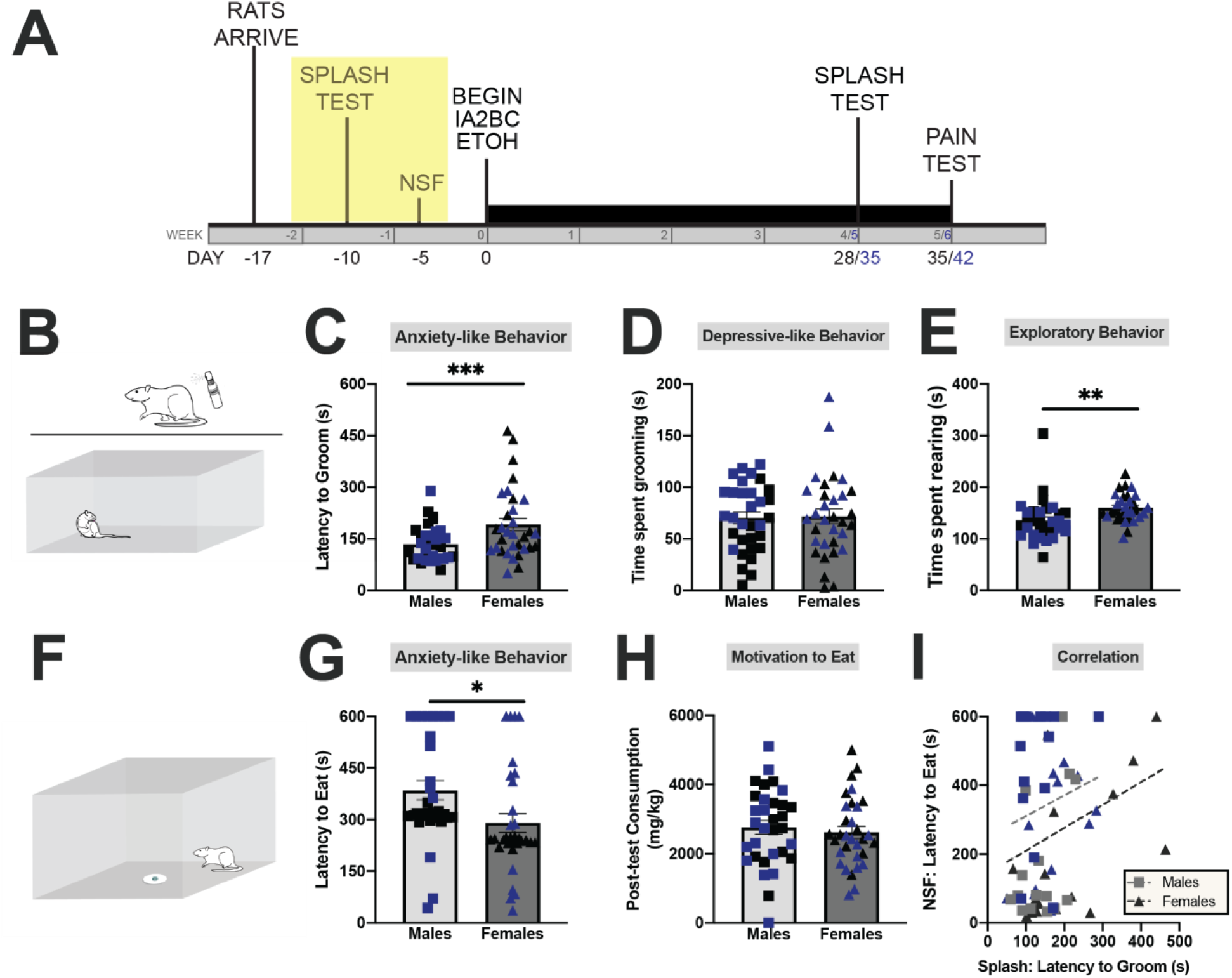
Sex differences in basal anxiety- and depressive-like behavior. **(A)** Experimental timeline of entire experiment. **(B)** Schematic of Splash Test. **(C)** Compared to males, females exhibit increased latency to groom. **(D)** No sex differences in time spent grooming. **(E)** Females spent more time rearing compared to males **(F)** Schematic of Novelty Suppressed Feeding Test. (**G**) Males exhibit increased latency to eat. **(H)** No sex differences in post-test consumption. (**I).** There is no significant correlation between behavior on the splash test and novelty suppressed feeding tests. Data expressed as individual points with bars representing mean + SEM. Separate cohorts are represented as black or blue. Panels C-E and G-H utilized an unpaired student’s t test (N=32/sex) whereas panel I used Pearson’s correlation. *p<0.05, **p<0.01, ***p<0.001.

### Voluntary Ethanol Drinking

Following baseline behavioral testing, rats were allowed to voluntarily drink 20% ethanol (EtOH) and water (**Fig 2A**) under an intermittent access two bottle choice drinking paradigm (**Fig 2B**). We found no sex differences in EtOH intake when data were analyzed by individual drinking session (**Fig 2C**; Sex x Session: F(14, 416)=1.485; Sex: F(1,30)=3.61e-005; Session: F(14,416)=2.336, p<0.01) or when EtOH intake was averaged by week (**Fig 2D**; Sex x Session: F(4,120)=2.230, Sex: F(1,30)=0.001, Session: F(4, 120)=1.885). During the 5^th^ week of EtOH drinking, we measured intake at 1h and found no sex differences (**Fig 2E;** Session x Sex: F(2,60)=0.720, Sex: F(1,30)=0.105, Session: F(2,60)=2.756). Interestingly, both male and female rats drink approximately half of their total 24 hr EtOH intake in the first hour of access. Individual data points for **Fig 2C-2E** are shown in **2F-2H**, respectively, to illustrate group averages as well as individual variability in each measure.

**Figure 2.**
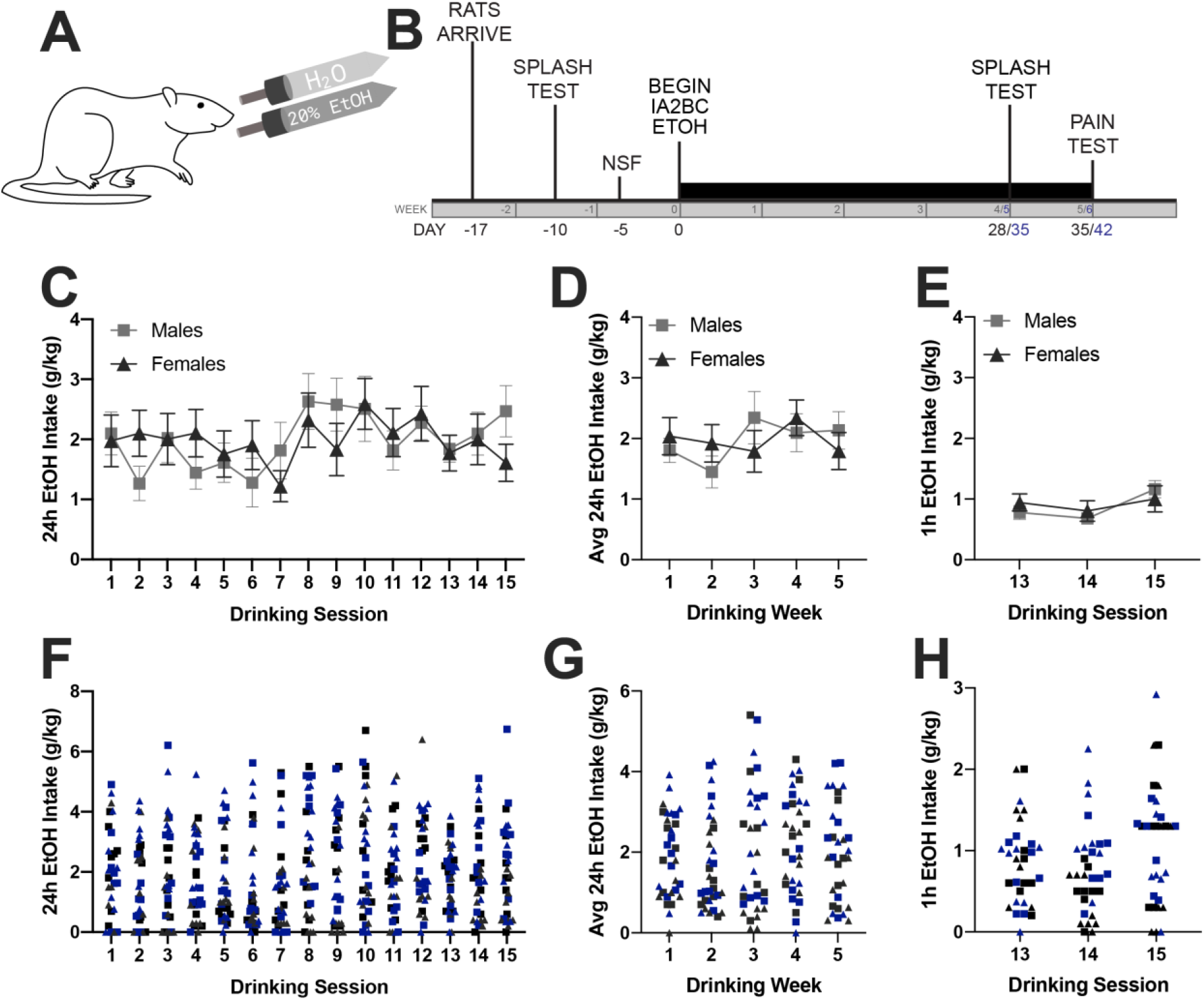
Voluntary EtOH Drinking Under an Intermittent Access Paradigm. **(A).** Schematic of drinking paradigm. **(B).** Experimental timeline. There are no effects of sex on 24h intake when analyzed by day **(C)** or week **(D).** There are also no effects of sex in 1h intake (**E**). Individual data points for (**C-E**) are shown in (**F-H**), respectively. Data are expressed as mean + SEM (**C-E**) or individual points (**F-H**). Separate cohorts are represented as black or blue in (**F-H**). N=16/sex.

### Baseline Affect and Subsequent Drinking

#### Anxiety-like Behavior

To examine if basal anxiety-like behavior was associated with EtOH intake, we performed correlational analyses on the anxiety-like behavior seen in the splash test and NSF with the first two weeks of subsequent EtOH intake. The experimental timeline with specific parameters being correlated are shown in **Fig S1A.** There was no significant correlation between either splash test (**Fig S1B**; males: r(14)=-0.140, females: r(14)=-0.411) or NSF (**Fig S1C**; males: r(14)=-0.1014, females: r(14)=-0.329) during the first week. In addition, the lack of association between anxiety-like behavior and drinking persisted into week 2 in both splash test (**Fig S1D**; males: r(14)=-0.097, females: r(14)=-0.257) and NSF (**Fig S1E**; males: r(14)=-0.170, females: r(14)=-0.214).

#### Depressive-like Behavior

The Splash test allows us to also examine depressive-like behavior in the total time spent grooming and the association of this behavior with drinking. Similar to above, these correlations were performed using the baseline splash test results and the first two weeks of drinking (**Fig S2A**). While basal depressive-like behavior was not significantly associated with drinking in the first week (**Fig S2B**; males: r(14)=-0.240, females: r(14)=0.432), a negative correlation emerged during week 2 of drinking (**Fig S2C;** males: r(14)=-0.357, females: r(14)=0.550, p<0.05). Interestingly, when examining the week 2 correlations, female rats exhibiting increased depressive-like behavior (i.e. decreased total time spent grooming) drank less EtOH while there was no relationship between depressive-like behavior and drinking in males.

#### Exploratory Behavior

Lastly, we correlated exploratory behavior in the splash test with drinking during the first 2 weeks (**Fig S3A**). While there were sex differences in basal exploratory behavior, there were no correlations of rearing behavior with drinking during week 1 (**Fig S3B**; males: r(14)=0.400; females: r(14)=-0.106) or week 2 (**Fig S3C**; males: r(14)=0.338,females: r(14)=-0.292).

### Negative Affect After Voluntary Drinking

Because of the correlation of depressive-like behavior and voluntary drinking, we wanted to investigate if a history of EtOH drinking could alter subsequent negative affect. Thus, we repeated the splash test in all animals (EtOH drinking and water controls) and compared the behavior to the baseline test (**Fig 3A**). We found no effect of chronic EtOH drinking on anxiety-like behavior in males (**Fig 3B**; EtOH x Time: F(1,30)=0.015, EtOH: F(1,30)=1.670, Time: F(1,30)=9.884e-005) or females (**Fig 3C**; EtOH x Time: F(1,29)=0.357, EtOH: F(1,29)=1.664, Time: F(1,29)=0.368). However, there was an effect on depressive-like behavior in males, where the total time spent grooming was significantly decreased in rats from both water and EtOH groups (**Fig 3D**; EtOH x Time: F(1,30)=0.001, EtOH: F(1,30)=0.033, Time: F(1,30)=12.30, p<0.01). Interestingly, this effect was absent in females (**Fig 3E**; EtOH x Time: F(1,29)=4.018, EtOH: F(1,29)=0.163, Time: F(1,29)=0.160). Lastly, there was no effect of EtOH drinking on exploratory behavior in either males (**Fig 3F**; Males: EtOH x Time: F(1,30)=0.699, EtOH: F(1,30)=0.123, Time: F(1,30)=1.330) or females (**Fig 3G**; EtOH x Time: F(1,29)=4.121, EtOH: F(1,29)=0.005, Time: F(1,29)=2.002). Together, these data suggest that chronic EtOH drinking does not alter negative affect as measured through the splash test.

**Figure 3.**
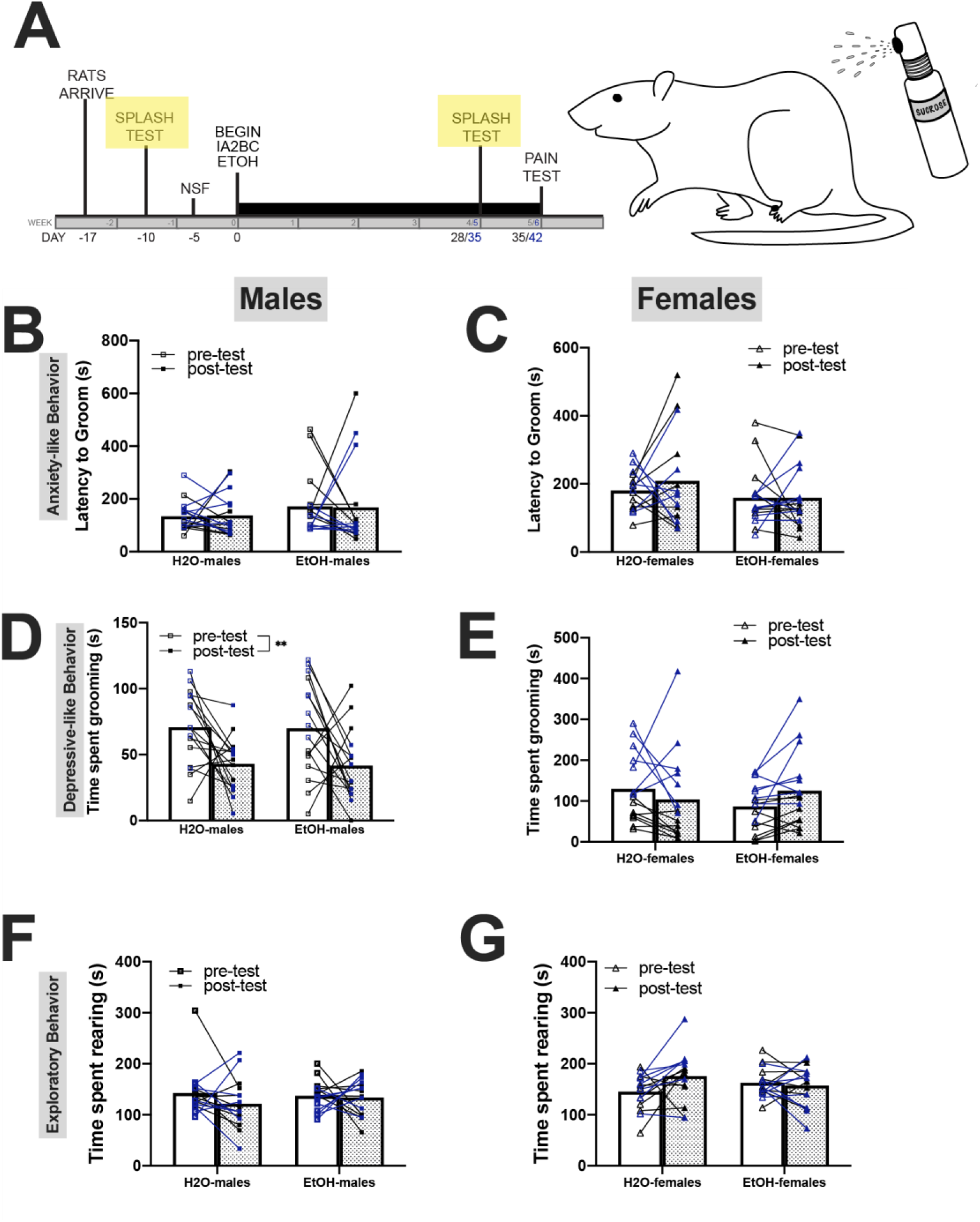
Negative Affect After Voluntary Drinking. **(A).** Experimental timeline highlighting pre- and post-tests. There is no effect of EtOH on latency to groom in males **(B)** or females **(C).** Similarly, there is no effect of EtOH on time spent grooming in males **(D)** or females **(E).** However, retesting results in decreased time spent grooming (2 way ANOVA) in males **(D)** but not females **(E).** Lastly, there was no effect of EtOH or re-testing in exploratory behavior in males **(F)** or females **(G).** Data expressed as individual points with bars representing mean. Separate cohorts are represented as black or blue. Data were analyzed with a 2 way ANOVA (N=15-16/group). **p<0.01

### Previous Drinking and Negative Affect

While a history of EtOH drinking did not affect subsequent anxiety- and depressive-like behavior, we wanted to further examine if there were any dose-dependent effects of EtOH i.e., if higher drinking levels correspond to increases in depressive-like behavior. Thus, we compared both the previous week and previous day EtOH intake to behavior in the splash test, as visualized in **Fig S4A**. We found no associations between previous week EtOH intake and anxiety-like behavior (**Fig S4B**; males: r(14)=-0.005432, females: r(14)=0.090), depressive-like behavior (**Fig S4C**; males: r(14)=0.0274, females: r(14)=-0.093), or exploratory behavior (**Fig S4D**; males: r(14)=-0.195, females: r(14)=0.317). Additionally, there were no dose-dependent effects when examining the association between the previous day’s EtOH intake and anxiety-like behavior (**Fig S4E**; males: r(14)=-0.011, females: r(14)=0.110), depressive-like behavior (**Fig S4F**; males: r(14)=0.064, females: r(14)=0.189) or exploratory behavior (males: r(14)=-0.025, females: r(14)=-0.141). These data suggest that prior EtOH drinking levels are not predictive of subsequent negative affect behavior.

### Associations between Negative Affect and Subsequent Drinking

As we previously observed that depressive-like behavior was negatively correlated with future EtOH intake in females, we wanted to examine if this correlation persisted after a history of chronic drinking. Thus, similar to analyses conducted in Figures S1-3, we performed Pearson’s correlations on the association between negative affect and subsequent drinking (as depicted in **Fig 4A**). In contrast to what we observed with baseline testing, we found that females that exhibited increased anxiety-like behavior drank significantly more EtOH during their next drinking session (**Fig 4B;** r(14)=0.679, p<0.01). Furthermore, this effect was specific to females, as there was no statistically significant correlation of anxiety-like behavior and EtOH intake in males (**Fig 4B**; r(14)=0.337). There was no association between depressive-like behavior (**Fig 4C**; males: r(14)=-0.0902, females: r(14)=-0.353) or exploratory behavior in either males or females **Fig 4E;** males: r(14)=0.220, females: r(14)=0.142). The positive correlation between anxiety-like behavior and drinking persisted into the following week in females (**Fig 4E**: r(14)=0.497, p<0.05). Consistent with the next day findings, we observed no correlation between anxiety-like behavior and drinking in males (**Fig 4E**: r(14)=0.164). There was also no association between total time spent grooming and drinking over the next week in either sex **(Fig 4F:** males: r(14)=-0.094, females: r(14)=-0.026). Similarly, there was no association between exploratory behavior and drinking over the next behavior **(Fig 4G:** males: r(14)=0.028, females: r(14)=0.145). Together, these data indicate that anxiety-like behavior, but not depressive-like or exploratory behavior, is associated with increased subsequent EtOH drinking in a sex-dependent manner.

**Figure 4.**
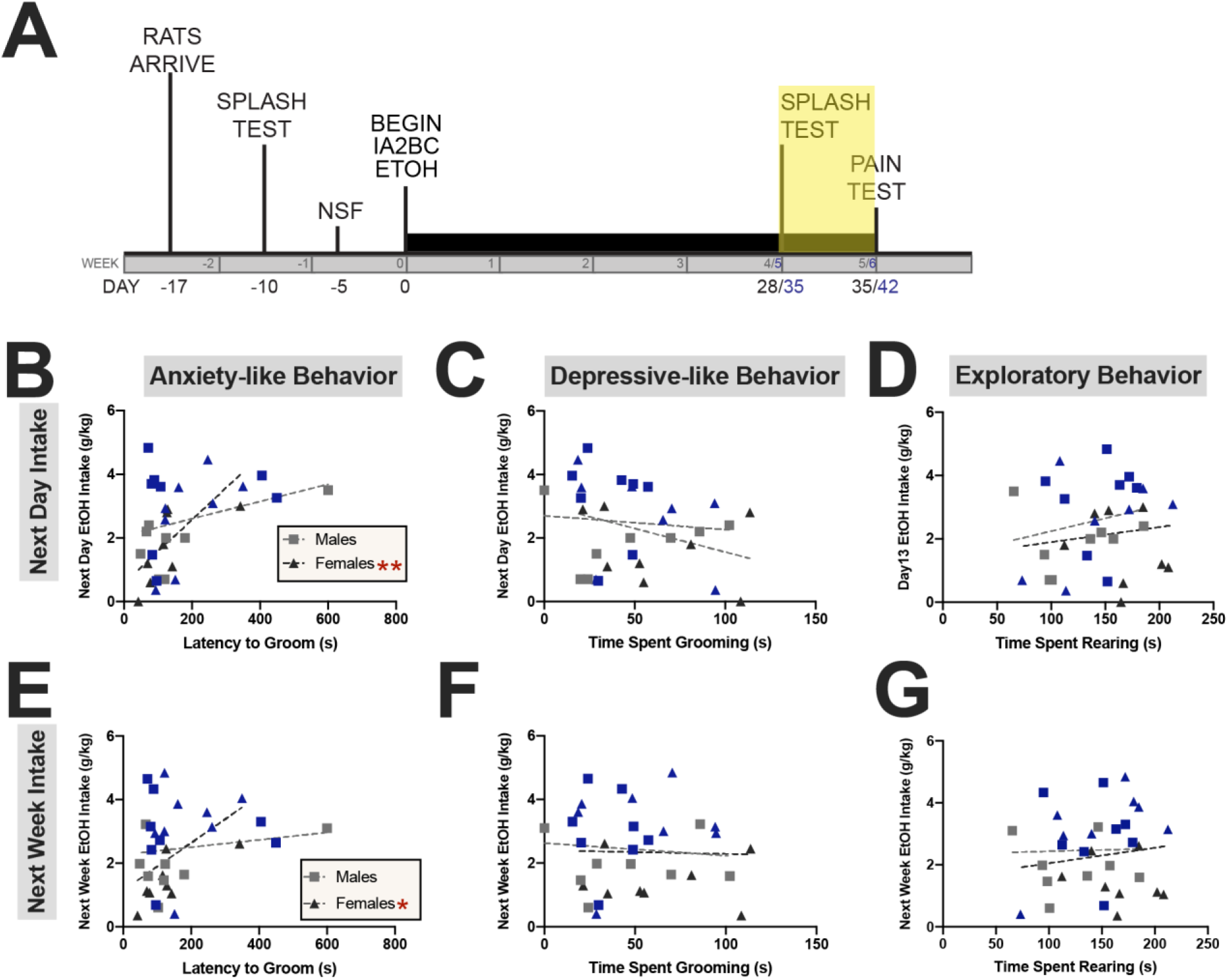
Associations between Negative Affect and Subsequent Drinking. **(A).** Experimental timeline highlighting behavioral testing and drinking to be correlated in subsequent panels. **(B)** Latency to groom was correlated with next day intake in only females. Time spent grooming **(C)** and time spent rearing **(D)** do not correlated with next day intake. **(E)** The correlation of latency to groom and future intake in females persists into the next week but is not correlated in males. Time spent grooming (F) and time spent rearing **(G)** do not correlate with next week drinking. Data expressed as individual points with lines of best fit. Separate cohorts are represented as black or blue. Data were analyzed with a Pearson’s correlation. *p<0.05 **p<0.01

### Thermal Sensitivity Following Chronic EtOH Drinking

While negative affect generally includes anxiety- and depressive-like behaviors, EtOH is also known to affect pain sensitivity. Using the Hargreaves apparatus (depicted in **Fig 5a**), we measured sensitivity to a thermal stimulus. We found that a chronic history of EtOH intake resulted in a decreased latency to withdraw in both males and females, indicative of a heightened pain response in both sexes (**Fig 5b**; Sex x EtOH: F(1,60)=0.0006, Sex: F(1,60)=2.818, EtOH: F(1,60)=6.165, p<0.05). However, in both males and females, thermal sensitivity was not correlated with previous day 1h ethanol intake (**Fig 5c**; males: r(14)=-0.137, females: r(14)=0.111), previous day 24h ethanol intake (**Fig 5d**; males: r(14)=-0.098, females: r(14)=0.149), previous week 1h ethanol intake (**Fig 5e**; males: r(14)=-0.102, females: r(14)=0.140) or previous week 24h intake (**Fig 5f**; males: r(14)=0.019, females: r(14)=-0.183) These data suggest that while chronic EtOH drinking results in increased sensitivity to thermal stimuli, this effect is not sex-specific or dose-dependent.

**Figure 5.**
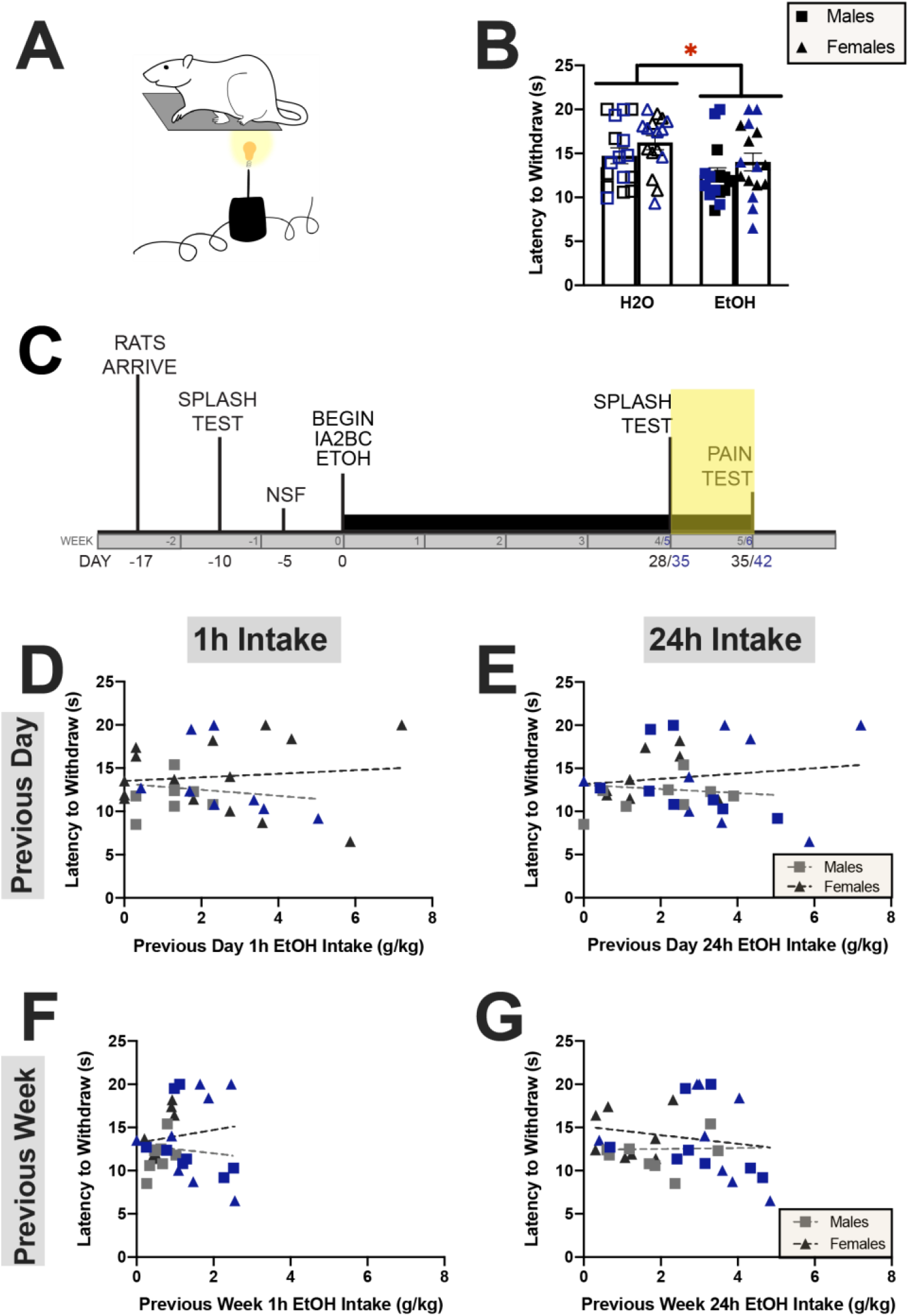
Thermal Sensitivity Following Chronic EtOH Drinking. (**A).** Schematic of Hargreaves test of thermal sensitivity**. (B)** Chronic EtOH intake increases thermal sensitivity in both male and female rats (2-way ANOVA). **(C)** Experimental timeline highlighting time of drinking and behavioral tests that are correlated in subsequent panels. Previous day (D-E) or week (F-G) EtOH is not associated with thermal sensitivity (Pearson’s correlation). Data expressed as individual points with bars representing mean + SEM (B). Data expressed as individual points with lines of best fit (D-G). Separate cohorts are represented as black or blue. N=16/group. *p<0.05

## DISCUSSION

The goal of the present study was to investigate the relationship between negative affect and ethanol drinking with the important consideration of sex as a biological variable. Using complementary tests of anxiety- and depressive-like behavior, we found sex- and test-specific differences in basal affective behavior such that females displayed enhanced anxiety-like behavior in a grooming test and males displayed enhanced anxiety-like behavior in a food-motivated test paradigm. Although there were no sex differences in ethanol drinking and no correlation between anxiety-like behavior and subsequent EtOH intake, we did find that depressive-like behavior predicted future ethanol intake in female rats only. In addition, while we observed no differences in anxiety-like behavior after chronic drinking, we did observe an increase in depressive-like behavior in male rats in both the water and ethanol drinking groups. Ethanol drinking levels did not correlate with performance on anxiety- or depressive-like behavioral testing in either sex. However, anxiety-like behavior, but not depressive-like behavior did predict subsequent ethanol intake in female but not male rats. Lastly, we found that a history of alcohol intake decreased pain thresholds in male and female rats, but this effect is not influenced by sex or amount of EtOH intake. Together, these experiments provide important information on the complex interaction between negative affect and alcohol intake and how these two contexts reciprocally do, or do not, influence each other in a sex-specific manner.

Many different tests of anxiety- and depressive-like behavior exist in preclinical rodent models. Most tests of anxiety-like behavior measure the rodent’s willingness to enter an unsafe area or hesitancy to initiate naturalistic behavior. In the NSF, rodents are food deprived (in order to increase motivation to eat) and then placed in a brightly lit arena with the non-novel palatable food item in the center. Traditionally, latency to approach the food item is interpreted as anxiety-like behavior. In order to control for motivation to eat, post-test food consumption in the home cage is measured. Some might argue that decreases in the amount eaten during the post-test is indicative of anhedonic or depressive-like behavior; however, this is confounded significantly by the food deprivation. Food-deprivation impairs stimulus processing and increases locomotor activity and exploration (Baumeister et al., 1964) while impairing fear extinction and associated lateral amygdala plasticity (De la Casa, 2013) (Huang et al., 2016). Additionally, food deprivation increases corticosterone levels (Nowland et al., 2011, Duriez et al., 2020), which may artificially inflate levels of depressive-like behavior.

The splash test, on the other hand, involves spraying a viscous (typically sucrose) solution on the rodent’s back then placing the rodent in the testing arena. As grooming is an intrinsically-motivated behavior (and sucrose solutions have an inherently palatable taste), the rodent should, in theory, immediately begin grooming. However, if the rodent is exhibiting increased anxiety-like behavior, it may take longer to initiate grooming. The total time spent grooming, or number of grooming bouts, is often interpreted as a measure of depressive-like behavior, as it mimics a lack of self-care and hygienic apathy in humans. In the current study, the arena for the splash test was illuminated by red light (rather than bright white light) to minimize anxiety-like behavior and also provide a different context from the NSF.

Studies have found varying results with regards to sex differences in anxiety-like behavior. Some studies report that male mice exhibit increased latency to eat in the NSF (Jury et al., 2017, Bloch et al., 2020), whereas other studies using Wistar rats report the opposite, with females exhibiting increased latency to eat compared to males (De Oliveira Sergio et al., 2021). Additionally, other studies saw no sex differences in anxiety-like behavior in the NSF in both Sprague Dawley (Wright et al., 2019) and Wistar (Olivier et al., 2008) rats. In contrast to previous reports, we found that male Sprague Dawley rats exhibit increased basal anxiety-like behavior in the NSF. One potential explanation for this discrepancy is differences in housing conditions and sex-specific sensitivity to housing condition. In the present study our animals were single-housed for one week prior to testing (in preparation for drinking) whereas other studies used group-housed animals (Wright et al., 2019). Indeed, studies have found that social isolation increases anxiety-like behavior (tested via NSF) in males but not females (Oliver et al., 2020). The Splash test is another test used to measure anxiety-like behavior that provides information about anxiety-like behavior (latency to groom) as well as depressive-like behavior (total time spent grooming). To our knowledge, we are the first to compare male and female rodents in the Splash test and we found that females exhibit increased anxiety-like behavior. The fact that both NSF and Splash test demonstrate a robust effect of sex but in opposing directions suggests that sex differences in anxiety-like behavior are influenced by test modality. This is further supported by the fact that the correlation of anxiety-like behavior between the two tests was not statistically significant.

After establishing baseline sex differences in negative affect, a subset of the rats underwent chronic intermittent EtOH drinking. Using the model first described by Wise (Wise, 1973), rats were allowed 24h access to an EtOH and water bottle on alternating days in the home cage environment. Hwa and colleagues found the intermittency in this model increases EtOH intake compared to continuous access (Hwa et al., 2011). Previous studies have found that female Long Evans and Wistar rats drink more than male rats (Aguirre et al., 2020, Morales et al., 2015, Sluyter et al., 2000, Priddy et al., 2017, McNamara and Ito, 2021). Interestingly, we saw no sex differences in EtOH intake in our studies, despite one study reporting that female Sprague Dawley rats drank more than male Sprague Dawley rats (Li et al., 2019). However, consistent with our findings, one previous study reported that male and female CD rats (which are derived from Sprague Dawley rats) do not differ in EtOH consumption in this model (Schramm-Sapyta et al., 2014).

Using this model, we found that Sprague Dawley rats drank approximately 2g/kg EtOH during their drinking sessions. Others have found that male Sprague Dawley rats drink 2-4g/kg (Bito-Onon et al., 2011, Li et al., 2011); however, this may be due to differences in light cycle and timing of EtOH presentation. Rats in the previous study were on a reverse light cycle (where EtOH is presented during the dark cycle) and our rats were maintained on a normal light cycle (where EtOH is presented during the light cycle). Interestingly, we found that rats drink approximately 50% of their total EtOH intake during the first hour of EtOH presentation, even though EtOH was presented during the light cycle, a period of decreased overall activity. Others have examined EtOH intake at the 30min mark and found that Sprague Dawley rats consume 20-25% of their total intake during the first 30min (Li et al., 2010), similar to Long Evans rats consuming 10-30% of their EtOH in the first 30min (Carnicella et al., 2009, Morales et al., 2015). Considering none of the previous studies measured 1h intake, it is hard to compare them directly with our results. Our results add to the existing literature demonstrating that a majority of EtOH intake occurs within the first hour of EtOH presentation, despite this occurring during the light cycle. It is also worthwhile to note that the animals drank no water during the first hour, suggesting an alcohol-specific effect. This may model compulsive-like ethanol drinking behavior in humans, where individuals with AUD will consume alcohol at first availability and in inappropriate settings.

Many human studies have found direct relationships between anxiety and EtOH intake, where increased levels of anxiety correlate with increased EtOH drinking (Crum et al., 2013). We, however, found no correlation between basal anxiety-like behavior and subsequent drinking in the present study. However, after chronic EtOH drinking, anxiety-like behavior did predict subsequent drinking in females only. Thus, perhaps drinking changes anxiety-like behavior (albeit not significantly) and the resultant anxiety-like behavior contributes to future EtOH intake. This would be consistent with the self-medicating hypothesis, where individuals drink more EtOH to overcome their anxiety that resulted from previous drinking experience (Koob, 2003, Smith and Randall, 2012). Additionally, this is consistent with the allostatic model of AUD (Koob, 2003), where chronic EtOH drinking changes the allostatic set point and thus the increased allostatic load (manifesting as anxiety-like behavior) results in increased EtOH intake.

One crucial piece of knowledge gained from these studies is that anxiety-like behavior predicted future drinking in female, but not male, rats. There are limited preclinical studies that assess individual differences in drinking behavior as it relates to anxiety-like behavior. One study grouped male Wistar rats into “high anxiety phenotype” and “low anxiety phenotype” based on performance in the elevated zero maze before undergoing EtOH home cage drinking via saccharin fading (Sgobbi and Nobre, 2020). The researchers found that rats with a high anxiety phenotype consumed more adulterated EtOH than rats in the low anxiety phenotype (Sgobbi and Nobre, 2020). However, once the saccharin was fully faded out, there were no group differences, which is consistent with our results (Sgobbi and Nobre, 2020). A separate study found that anxiety-like behavior on the elevated plus maze was positively correlated with operant EtOH self-administration in Marchigian Sardinian alcohol-preferring (msP), but not Wistar rats (Domi et al., 2019). However, there was no correlation between anxiety-like behavior and persistence in EtOH-seeking, resistance to punishment, or motivation for EtOH in either the msP rats or the Wistar rats (Domi et al., 2019). Unfortunately, these studies were performed only in males (Domi et al., 2019), so it remains unclear how or if their results extend to female rats.

One interesting finding from the present study was the lack of effect of EtOH drinking on anxiety- and depressive-like behavior in the splash test. The splash test measures both anxiety-like (latency to groom) and depressive-like (time spent grooming) behavior. Only two other studies, to our knowledge, have utilized the splash test to examine negative affect after chronic EtOH intake. One study in mice found that males that underwent 4 weeks of drinking in the dark exhibited increased anxiety-like behavior (latency to groom) and depressive-like behavior (time spent grooming) compared to water drinking controls (Sampedro-Piquero et al., 2020). One reason for the differences in our findings, in addition to inherent species differences, is that the mice used in the previous study drank significantly more than the rats used in the present study. The mice drank approximately 6g/kg while the rats in this study drank 2g/kg. Thus, the drinking levels in our study may not have met the threshold for producing changes in anxiety-like behavior. Another study using the splash test found no effect of intermittent EtOH drinking or chronic intermittent ethanol exposure on number of grooming bouts in mice, but failed to report any effect of sex or other relevant outcomes such as latency to groom and time spent grooming (Okhuarobo et al., 2020).

Our studies also found that 6 weeks of social isolation induces negative affect in male, but not female, rats. Many researchers have examined the effects of social isolation (single-housing) on anxiety-like behavior with mixed results, potentially due to the different tests of anxiety-like behavior used. Studies using the NSF as a readout of anxiety-like behavior found that social isolation increased anxiety-like behavior in male but not female mice (Oliver et al., 2020) whereas other studies found that social isolation increases anxiety-like behavior in the light-dark test and the open field test, but found no effect of social isolation using the elevated plus maze (Evans et al., 2020). We found no differences in anxiety-like behavior in both males and females following 5-6 weeks voluntary drinking during social isolation as measured by the latency to initiate grooming behaviors. Depressive-like behavior was measured as time spent grooming during the splash test. In this study, we found that social isolation increased depressive-like behavior in males, without affecting females, consistent with other studies that examined the effect of social isolation on sucrose preference (a measure of anhedonia) in mice (Evans et al., 2020, Oliver et al., 2020). However, there are mixed results in examining the effects of social isolation on females – some studies find social isolation decreases anhedonia (Oliver et al., 2020), but others found it increases anhedonia (Evans et al., 2020). There is a gap in the literature regarding the effects of social deprivation during adulthood in rats – many studies focus on the pre-weaning period or adolescent socialization. Additional studies are required to understand the influence of adult social isolation on negative affect in male and female rodents.

We also demonstrated that EtOH intake of 2g/kg is sufficient to produce hyperalgesia, as measured through the Hargreaves test of thermal sensitivity. Our results are consistent with other studies that have reported heightened pain states as a result of chronic intermittent ethanol vapor, (Avegno et al., 2018, Edwards et al., 2012), ethanol-containing liquid diet (Dina et al., 2006), or voluntary drinking (Quadir et al., 2020b, Quadir et al., 2020a, Quadir et al., 2021). However, we did not see a dose-dependent effect of EtOH or an effect of sex on baseline or EtOH-induced changes in thermal sensitivity. Other studies in mice have also demonstrated a lack of dose-dependency on pain states during EtOH withdrawal (Quadir et al., 2021). The lack of effect of sex on hyperalgesia is also consistent with previous studies demonstrating that alcohol’s analgesic effects are not sex-specific (Bilbao et al., 2019, Neddenriep et al., 2019).

One major caveat of this study is that we did not monitor the estrous cycle of the female rats. As monitoring estrous cycle involves handling and manipulation that could not be properly controlled for in male rats, we chose not to specifically measure estrous cycle in the present study. While some tests of anxiety-like behavior are unaffected by the estrous cycle (elevated plus maze and open field, (Scholl et al., 2019)), other studies have found that females in the diestrous stage display increased anxiety-like behavior in the NSF (Mora et al., 1996). In addition, while others have not examined the effect of estrous cycle stage on grooming behaviors, others have found female rats in diestrous spend more time floating in the forced swim test (Marvan et al., 1996, Marvan et al., 1997, Consoli et al., 2005) and have higher escape latencies in tests of learned helplessness (Jenkins et al., 2001) suggesting estrous stage-dependent changes in depressive-like behavior. The role of the estrous cycle in mediating EtOH intake is unclear – some studies have found no effect (Ford et al., 2002, Priddy et al., 2017), but others have found that EtOH intake peaks during diestrus (Ford et al., 2002). Lastly, researchers have also found that mechanical sensitivity (Von Frey test), but not thermal sensitivity (Hargreaves test) or inflammation-induced pain (Formalin test) are mediated through by the estrous cycle (Yang et al., 2020). Future studies are required to clarify the role of the estrous cycle in EtOH drinking and negative affect.

As evident in both preclinical and clinical work, AUDs are extremely complex conditions that are intertwined with mood disorders such as anxiety and depression. Through the use of rodent models of voluntary ethanol drinking and behavioral testing, our studies provide important evidence for sex differences in basal anxiety- and depressive-like behaviors and their ability to predict subsequent EtOH-induced negative affect. Our studies also highlight the importance of examining individual differences, which will hopefully allow for a shift toward personalized medicine as a treatment for AUD.

## AUTHOR CONTRIBUTIONS

S.G.Q and M.A.H, designed and analyzed experiments. S.G.Q., G.M.A., J.V.J., and M.E.S. performed experiments. S.G.Q. and M.A.H. wrote the manuscript.

## ACKNOWLEDGEMENTS

This work was supported by National Institute of Health grants AA011605 (M.A.H), AA026858 (M.A.H) and AA007573 (S.G.Q). We thank Sara Y Conley for her assistance in pilot experiments.

## Notes

### Competing Interest Statement

The authors have declared no competing interest.

