## Supplemental Materials for "Sex Differences in Affective States and Association with Voluntary Ethanol Intake in Sprague Dawley Rats"

**Figure S1**

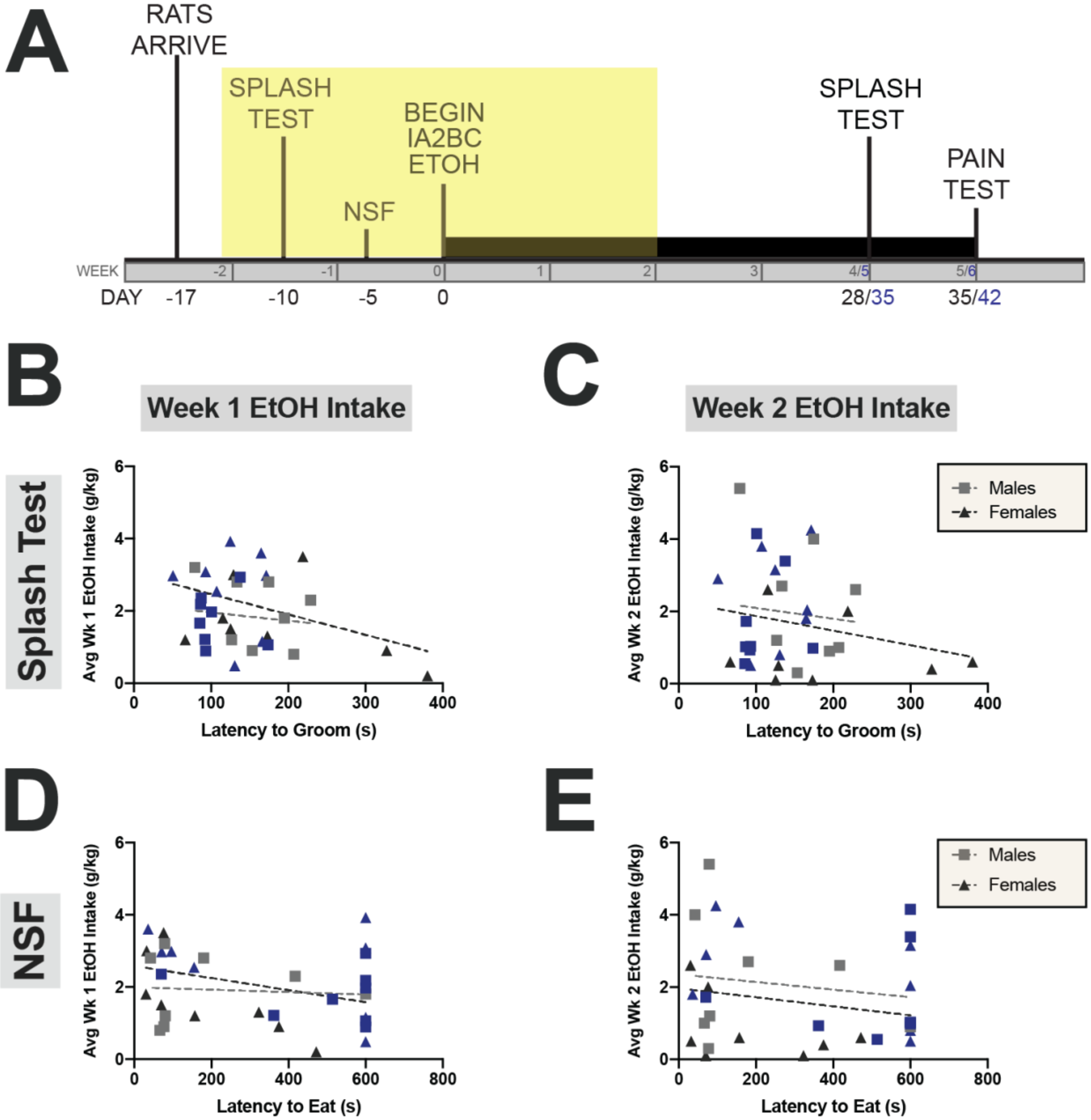


**Figure S1. Basal anxiety-like behavior is not associated with EtOH Intake**

**(A).** Experimental timeline highlighting behavioral testing and drinking to be correlated in subsequent panels. Latency to groom in the splash test does not predict week 1 **(B)** or week 2 **(C)** EtOH intake. Latency to eat in the NSF does not predict week 1 **(D)** or week 2 **(E)** intake.

Data expressed as individual points with lines of best fit. Separate cohorts are represented as black or blue. N=16/group.

**Figure S2**

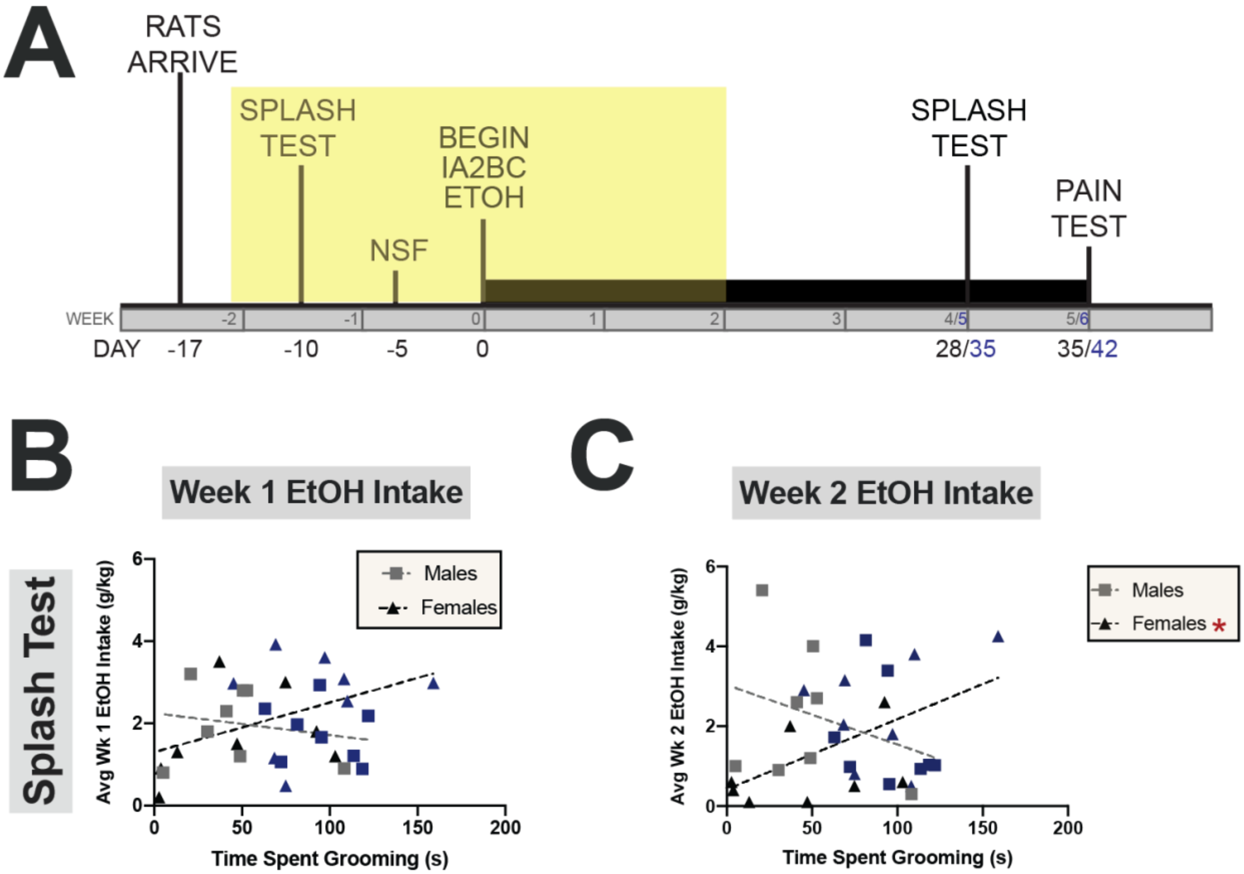


**Figure S2. Basal depressive-like behavior is associated with future EtOH in female but not male rats.**

**(A).** Experimental timeline highlighting behavioral testing and drinking to be correlated in subsequent panels. Time spent grooming does not predict week 1 intake in males or females **(B)** but does predict week 2 intake in females **(C).**

Data expressed as individual points with lines of best fit. Separate cohorts are represented as black or blue. N=16/group. *p<0.05 (Pearson's correlation)

### Figure S3

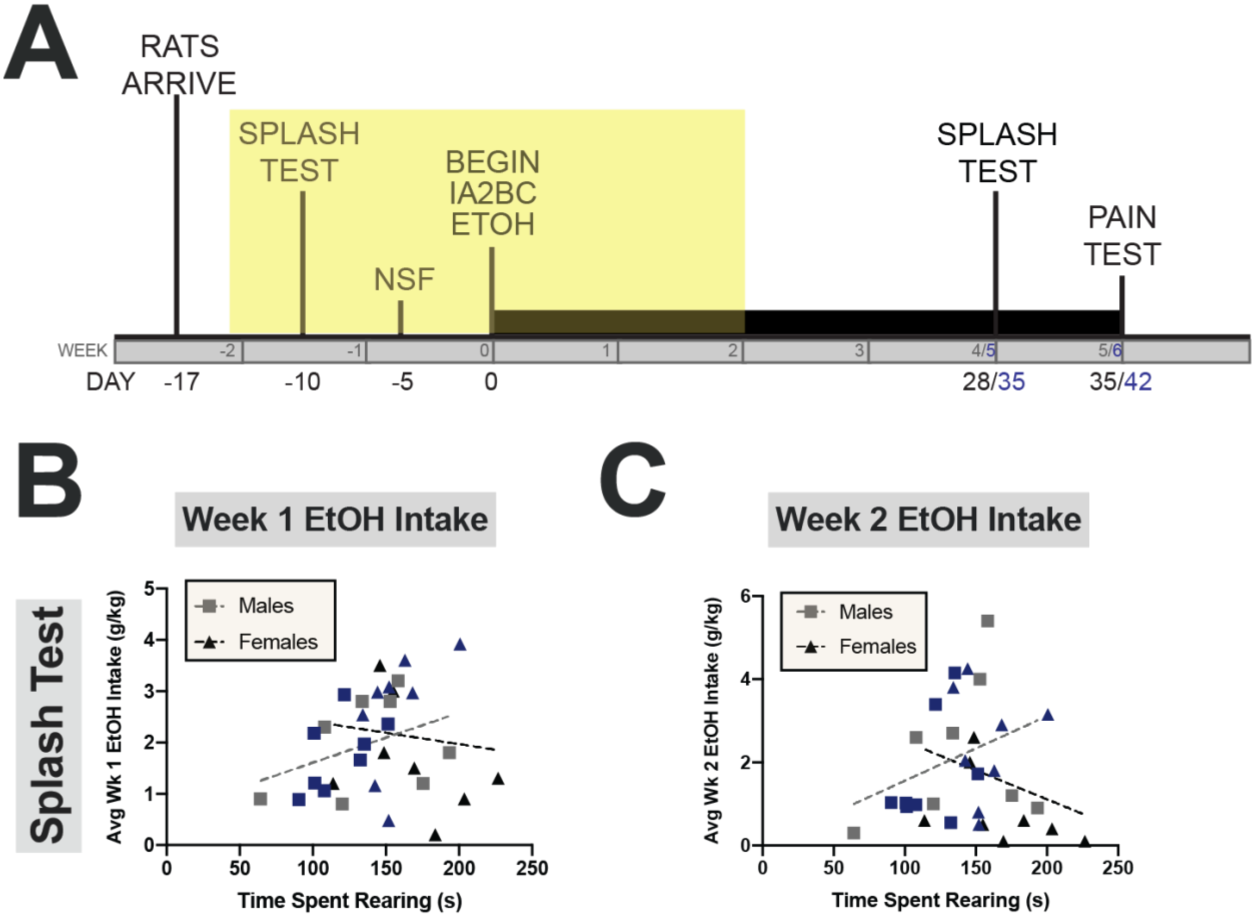


**Figure S3. Basal exploratory behavior is not associated with future EtOH Intake (A).** Experimental timeline highlighting behavioral testing and drinking to be correlated in subsequent panels. Time spent rearing in the splash test does not predict week 1 **(B)** or week 2 **(C)** intake.

Data expressed as individual points with lines of best fit. Separate cohorts are represented as black or blue. N=16/group.

**Figure S4**

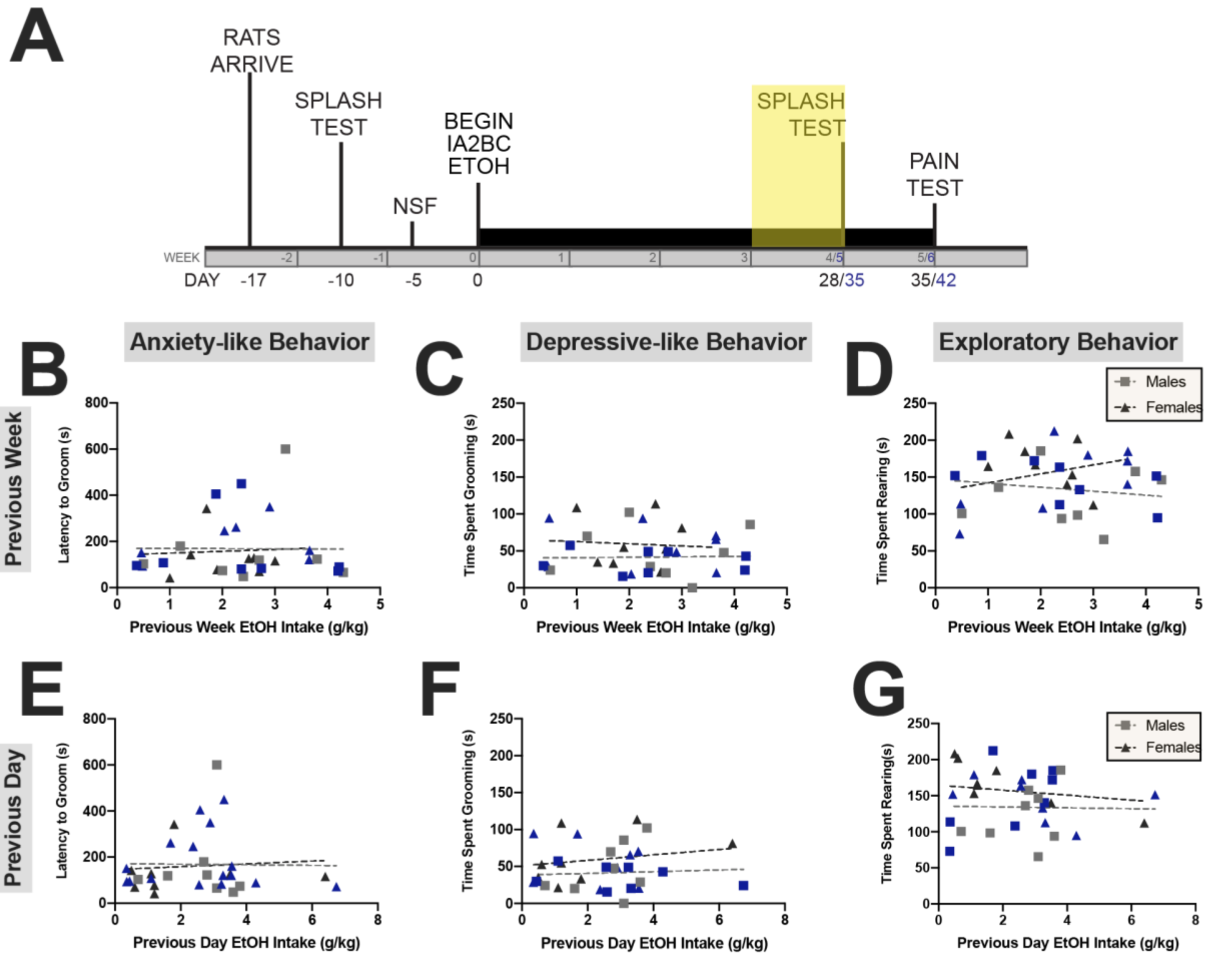


**Figure S4. Previous EtOH intake is not associated with affective state in the Splash Test**

**(A).** Experimental timeline highlighting drinking and behavioral testing to be correlated in subsequent panels. Previous week intake is not associated with latency to groom **(B),** time spent grooming **(C),** or time spent rearing **(D).** Similarly, previous day intake is not associated with latency to groom **(E),** time spent grooming **(F),** or time spent rearing **(G).**

Data expressed as individual points with lines of best fit. Separate cohorts are represented as black or blue. N=15-16/group.
